# Modeling meiotic chromosome pairing: increased fidelity from a tug of war between telomere forces and a pairing-based Brownian ratchet

**DOI:** 10.1101/332767

**Authors:** Wallace F. Marshall, Jennifer C. Fung

## Abstract

Meiotic homolog pairing involves associations between homologous DNA regions scattered along the length of a chromosome. When homologs associate, they tend to do so by a processive zippering processive, which apparently results from avidity effects. Using a computational model, we show that this avidity-driven processive zippering reduces the selectivity of pairing. When active random forces are applied to telomeres, this drop in selectivity is eliminated in a force-dependent manner. Further simulations suggest that active telomere forces are engaged in a tug-of-war against zippering, which can be interpreted as a Brownian ratchet with a stall force that depends on the dissociation constant of pairing. When perfectly homologous regions of high affinity compete with homeologous regions of lower affinity, the affinity difference can be amplified through this tug of war effect provided the telomere force acts in a range that is strong enough to oppose zippering of homeologs while still permitting zippering of correct homologs. The degree of unzippering depends on the radius of the nucleus, such that complete unzippering of homeologous regions can only take place if the nucleus is large enough to pull the two chromosomes completely apart. A picture of meiotic pairing thus emerges that is fundamentally mechanical in nature, possibly explaining the purpose of active telomere forces, increased nuclear diameter, and the presence of “Maverick” chromosomes in meiosis.

## Introduction

The pairing of homologous chromosomes in meiosis underlies all of Mendelian genetics, and yet its biophysical basis is poorly understood. Chromosomes are gigantic polymers, densely tangled inside the nucleus. How do homologous chromosomes search for the correct partner from amongst all the other potential pairing partners? Pairing requires that homologous DNA sequences be able to find each other and become physically associated, despite being initially separated at random locations on the order of microns apart. Two questions are how the large scale motions necessary for pairing are achieved, and how correct pairing partners are assessed during the search process. A key element of the first question is the possible role of active mechanical forces in catalyzing the motion of chromosomes through the densely tangled nucleus. A key element of the second question is how correct homologs are discriminated from incorrect pairing partners during the pairing process. Assessment of correct versus incorrect pairing partners is hypothesized to rely either on DNA-level sequence homology testing, requiring that transient base pairing occurs on a rapid time scale as the chromosomes move around, or else to rely on a chromosome-specific pattern of cohesins that would serve as a bar-code and allow homologs to recognize each other without the need for single-strand invasion and base pairing (Ishiguro et al., 2014). Another early theory for homolog recognition proposed proteins that recognize chromosome-specific sequences (Maguire 1984), a mechanism that is now known to operate in C. elegans (Phillips and Dernburg, 2006). As with all molecular recognition processes, the potential for errors in recognition always exists, raising the question of how meiotic pairing ensures that only the correct partner is ultimately recognized. These two key questions about pairing, namely the role of active forces in chromosome motion, and the mechanisms to ensure correct pairing, are usually thought to be distinct questions. Here we provide evidence, based on a computational modeling study, that in fact these two questions may be intertwined, such that active motion may play a role in correcting mistakes in pairing in addition to its role in speeding up initial pairing. The fact that motion could play both roles raises the question, for future research, of which role is actually more important.

Active chromosome motion has long been observed in meiosis, where it is thought to play a functional role in the process (Parvinen and Söderström, 1976; Sheehan and Pawlowski 2009). Meiotic chromosome motion is now known to be driven by actin-myosin or dynein-microtubules, depending on the species (Chikashige 1994; Scherthan 2007; Conrad 2008; Koszul 2008; Sato 2009). These cytoskeletal force generating units drive large random motions of telomeres, resulting from a coupling of telomeres to the cytoskeleton through the nuclear envelope via SUN and KASH domain proteins such as *csm4/mps3* in budding yeast (Conrad 2008; Koszul 2009; Sonntag Brown et al., 2011). Active motion in meiosis seems to be a highly conserved feature across different species, but what is this motion for? One hypothesis is that active motion helps to speed up the random search for homologous sequences in the crowded environment of the nucleus (Conrad 2008; Kosaka 2008; Mine-Hattab and Rothstein, 2012; Marshall and Fung, 2016). Mutations affecting telomere coupling to actin reduce the rate of collision between homologous loci (Lui et al., 2013) but also show actin-independent effects. Motion impairment delays meiotic completion (Rao 2011), but can increase the overall number of crossovers (Kosaka 2008; Wanat 2008). Transient arrest of active motion in fission yeast meiosis indicates that loss of motion delays initial pairing kinetics but also results in hyperstable pairing associations accompanied by unresolvable recombination events that cause a failure of chromosome segregation (Chacon et al., 2016). These complicated phenotypes raise the possibility that meiotic motion might play other roles beyond just facilitating the collision of chromosomes.

An alternative idea is that active forces might contribute to proper pairing by actively pulling apart weakly associated interactions. A precedent for this general concept that active forces can increase fidelity of a molecular pairing interaction has already been set by experimental studies of B Cell Receptor (BCR) antigen affinity discrimination (Natkanski et al., 2013). The interaction between the BCR and its antigen ligand is subject to mechanical tension generated by myosin IIa moving on actin, and this active pulling force plays a role in antigen affinity discrimination. Low affinity antigen-BCR interactions are ruptured by the myosin-generated force, such that only high affinity antigens can be stably captured by the receptor. This work demonstrates that mechanical forces generated by the cytoskeleton are able to enforce discrimination of molecular interactions. This mechanical testing is thought to be necessary because the BCR finds ligands with such a low off-rate that even a low affinity interaction (by BCR-antigen standards) still is strong enough to resist disruption by collisions with thermally excited solvent molecules. By adding a larger energetic contribution, active forces allow low-affinity interactions to be disrupted, thus providing discrimination that would be impossible by thermal energy alone at ordinary temperatures.

In addition to the studies of B cell receptor, mathematical and biophysics studies of single molecule dissociation provide a theoretical basis for the idea that force can probe the strength of molecular interactions. Bell’s formula (Bell 1978) predicts that the probability of two interacting molecules dissociating in any given time interval is an exponential function of the force pulling the objects apart, a prediction confirmed in force-mediated dissociation of cell-cell adhesions (Evans et al., 1991). Similar considerations have been used to develop methods for extracting dissociation rate constants from single molecule pulling experiments (Hummer and Szabo 2003). The ability of mechanical forces to probe the binding affinity of a molecular interaction is thus well established in other areas of biology and biophysics.

We hypothesize that cytoskeletal forces applied to telomeres in meiosis might play an affinity discrimination function during homology searching, similar to the role of myosin generated forces at the B cell receptor, by helping to erase incorrect associations stable enough to resist thermal dissociation. The need for mechanisms to ensure meiotic fidelity is indicated by studies showing that meiotic recombination can occur not only between allelic positions on homologs, but also between homologous sequences dispersed throughout the genome (Lichten et al., 1987; Montgomery 1991; Goldman 1996; Jinks-Robertson 1997), a phenomenon known as ectopic recombination. Ectopic recombination leads to incorrect chromosome segregation and meiotic defects. Given the prevalence of whole and partial genome duplications during evolution, the potential is high for partially homologous sequences to exist in multiple places in any given genome, which, if they recombined with each other, could lead to chromosome rearrangements or overt meiotic failure. Several mechanisms have been proposed to help prevent ectopic recombination, including masking of highly repeated regions by chromatin compaction and progressive alignment of homologs via pairing prior to onset of actual recombination. Could mechanical forces generated by actin/myosin pulling on telomeres also play an important role in opposing ectopic recombination by destabilizing interactions between partially homologous regions of the genome? If active force could pull apart weakly paired sequences, this would favor recombination between true homologous regions that share the highest sequence identity and are thus, presumably, more stably paired and hence better able to resist being pulled apart.

The affinity of two DNA sequences is a function of the level of homology, so that partially homologous regions anneal with less affinity than perfect homologs. But depending on the off-rate for dissociation of DNA strands, thermal motion might not be strong enough to drive dissociation of even partially homologous regions. This is especially true in light of the phenomenon of zippering. Ultrastructural studies of meiotic chromosomes have observed that pairing and synapsis occur through the elongation of a few initially paired tracts, rather than many simultaneous contacts. This so-called “zippering” is explained as an avidity phenomenon, whereby the pairing of a given locus constrains neighboring loci to be near each other, thus increasing their likelihood of colliding and pairing. Zippering has long been recognized as a key process in meiotic chromosome association (Zickler, 1972). Zippering is thought to account for the classic observation of Y-shaped chromosome arrangements during meiosis, in which homologs are paired starting at one end and continuing over part of their length, with the remainder of the homologs unpaired, a configuration known as “amphitene” (for review of this literature, see Wilson 1928). In our previously published simulations of meiotic pairing (Marshall and Fung, 2016) we found that even low-affinity pairing interactions, as reflected in a high probability of unpairing, led to processive pairing of the whole chromosome. The ability of low affinity interactions to drive processive pairing creates a potential problem for achieving high fidelity of pairing – if even poorly matched loci can drive processive zippering, how can correct interactions be distinguished from incorrect?

In the present study, we use simulations to investigate how zippering affects the ability to discriminate correct from incorrect pairing, as might happen in the presence of homeologous chromosomes, and whether active forces applied at telomeres might be able to restore fidelity of pairing. We find that active forces can indeed promote correct pairing, and we suggest that this effect can be understood as in terms of a Brownian ratchet mechanism.

## Methods

### Modeling framework for meiotic chromosome motion and pairing

We modeled meiotic chromosome dynamics using a previously described Brownian dynamics model. Each chromosome is represented as a list of nodes representing beads connected by springs, with each node subjected to Langevin random forces. The frictional force acting on each node is proportional to its velocity. These forces yield the following equation of motion, which is used to update the velocity and position of each node at each time step:

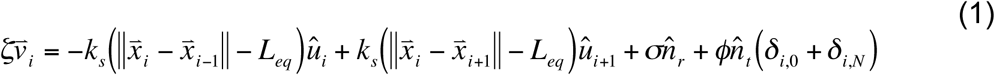

where x_i_ and v_i_ are the position and velocity vectors at node i, ζ is the friction coefficient, *k*_s_ and *L*_eq_ are the spring constant and equilibrium length of the links, σ is the magnitude of the Langevin (thermal) random force, ϕ is the magnitude of the cytoskeletally generated random force acting at the telomeres, n_r_ and n_t_ are randomly oriented unit vectors, and u_i_ represents a unit vector directed from node i-1 to node i. In the current version of the model, successive random oriented unit vectors are completely independent of each other, such that both the Langevin and telomere forces fluctuate in direction. We note that the term describing the Langevin random force actually represents the average net resultant of all random collisions with thermally excited solvent molecules during a single time-step. In our simulations the random unit vectors describing the motion of the telomeres are taken to always be tangential to the surface of the nucleus. We note that our representation of telomere motion, as a set of randomly oriented forces acting tangential to the nuclear surface and uncorrelated with the telomere forces on other chromosomes, would not be applicable in fission yeast, which differs from other systems in that the telomeres remain closely juxtaposed during the active movements and active forces drive large motions of the whole nucleus.

This bead-spring chain model represents a Rouse model of polymer dynamics (Grosberg and Khokhlov 1994) and thus assumes freely rotating links and lack of a barrier to strand passage. Such Rouse models of polymer dynamics have been shown to be consistent with experimental measurements of interphase chromatin motion (Hajjoul et al., 2013). The Rouse model is a “phantom polymer” model that lacks topological constraints. Order of magnitude scaling arguments based on the relative density of topoisomerase strand-passage sites compared to the density of chromatin interlocks have shown that there is effectively no barrier to strand passage because whenever an interlock would form, it is resolved by topoisomerase activity on a sufficiently rapid timescale (Sikorav and Jannink, 1994). Experimental studies have confirmed that topoisomerase activity can convert an entangled DNA melt into a viscous fluid, indicating that strand passage can readily occur on a times-scale relevant to chromosome motion (Kundukad and van der Maarel, 2010). Topoisomerase II activity is observed during meiosis, and defects in topoisomerase II activity lead to rapid arrest of meiosis (Rose and Hold, 1993; Zhang et al., 2014), confirming that this enzyme is active in strand-passage during the stage we are simulating. Obviously an interesting area for future computational analysis will be the effect of limitations on strand-passage rates. We note that one advantage of using the Rouse model is that our results could in principle be linked to analytical theory of polymers in a straightforward manner.

Initial generation of a random walk chain and relaxation according to equation 1 have been previously described (Marshall and Fung, 2016). Unless otherwise specified, all simulations in this paper run for 100000 time steps. The chromosomes are confined inside a spherical nuclear envelope by a repulsive force applied to any node moving outside the spherical shell in a direction normal to the surface with a spring constant k_nuc as previously described (Marshall and Fung, 2016). Telomeres are constrained to be located on the nuclear surface by calculating the radial distance from each telomeric node and then translating the telomere along the radius by this distance, so that after each step the telomere is always returned to a position on the nuclear surface (Marshall and Fung, 2016).

Pairing is modeled on an individual node basis, such that node 1 on a chromosome is only allowed to pair with node 1 on the homolog, node 2 with node 2, and so on. The simulations in this paper assume that all nodes are able to undergo pairing, although this is an adjustable parameter. As the homologous chains undergo their random movement, whenever corresponding loci move with a defined capture radius of each other, and if the local orientation of the two chains is closer to parallel than to antiparallel as judged by the dot product of a local orientation vector, they are set to a paired state and thereafter forced to move together until such time as they become unpaired. Unpairing of paired loci occurs with a uniform probability P_unpair per iteration. Affinity between a given pair of loci is reflected by the value of P_unpair, such that tightly associating loci (representing correct homologous associations) will have a low value for P_unpair, while incorrect associations would have a higher value for P_unpair. Thus, in our model, discrimination between correct and incorrect associations is reflected entirely by a difference in off-rates.

Table 1 gives the parameter values used for the simulations presented here, following order of magnitude estimates previously described (Marshall and Fung, 2016). Our model is not intended to precisely represent the actual chromosomes of any particular species, but rather is designed to be an abstract model that captures essential features of meiotic chromosome pairing. For this reason, our primary concern in choosing parameters is only to ensure that they are of the correct relative orders of magnitude.

**Table 1.** Simulation parameter values

| Model Parameter | Variable name | Parameter value | Physical units |
| --- | --- | --- | --- |
| Number of nodes | n_nodes | 50 | 500 kb |
| Frictional coefficient | $\zeta$ | 1.0 | $4 \times 10^{-6}$ Kg/s |
| Equilibrium spring length | $L_{eq}$ | 1.0 | 100 nm |
| Spring constant | $k_s$ | 0.25 | 0.007 pN/nm |
| Nuclear radius | nuc_radius | 5 | 0.5 μm |
| Repulsive force constant for NE | k_nuc | 0.35 | 1 pN |
| Capture radius for pairing | dist_capture | 0.5 | 50 nm |
| Magnitude of Langevin random force | $\sigma$ | 0.15 | 0.4 pN |
| Magnitude of telomere force | $\phi$ | 0.15–0.6 | 0.4 – 1.6 pN |
| Unpairing probability | P_unpair | 0.001-0.1 | 0.01 – 1.0 s<sup>-1</sup> |

Briefly, we employ a length scale in which 1 unit of length in the simulation corresponds with 100 nm of length in an actual cell. The equilibrium spring length for the links between nodes was chosen to be 1 unit (100 nm), consistent with the reported persistence length of yeast chromatin (Dekker, 2008). With this length scale, a nuclear radius of 5 units corresponds to a nuclear diameter of 1 micron, as is appropriate for yeast nuclei. Assuming that each link of 1 unit (100 nm) corresponds to 10 kb, based on prior estimates setting the packing density of chromatin at 110-150 bp/nm (Bystricky et al., 2004), a 50 node chain would correspond to a fully stretched chromosome length of 500 kb, which is in the range of actual yeast chromosome size. Capture distance was chosen based on the picture of a chromosome chain as a series of random polymer globules, and assuming that the diameter of each such globule equals the statistical link length. Based on the physical model of a polymer as a chain of random globules each corresponding to a statistical segment with diameter equal to the link length, this choice of capture distance corresponds to pairing taking place when the distance between the centers of the random globules of two homologous loci are separated by their radii, thus indicating a substantial degree of overlap between the two globules.

We set a timescale of 100 ms per simulation time step. With this assumption, our simulation of 100,000 time steps corresponds to a pairing time on the order of 5 hours, consistent with observations in yeast (note here we refer specifically to pairing and not to subsequent events of meiosis such as synapsis or crossing-over). Based on measurements of the diffusion constant (500 nm^2^/s) for yeast chromatin (Marshall et al., 1997) using the relation δ=sqrt(2ΔtD), we obtain a root mean-squared (rms) displacement during a single time step of δ=10 nm. This displacement is produced by thermal energy of a magnitude k_B_T, which we take as approximately 4 pN nm inside a yeast cell grown at 30C. Hence the rms force σ acting to drive this displacement is k_B_T/δ or 0.4 pN.

To derive the frictional coefficient ζ, we note that a node of the chain moves by δ=10 nm in a single timestep corresponding to 0.1 s, creating an apparent velocity of 100 nm/s. From the relation ζ=σ/v we obtain a value for of 4×10^−3^ pNs/nm which is equivalent to 4×10^−6^ Kg/s. To see if this is reasonable, we can calculate the viscosity corresponding to this frictional coefficient assuming Stoke’s law for a perfect sphere, such that η= ζ/6πR. Assuming the de Gennes model of spherical globules whose diameter matches the link length, we take R=50 nm, which yields a value for viscosity of 4 Pa s. This is approximately 4000 times the viscosity of pure water. The viscosity for the fluid phase of nucleoplasm, experienced by protein-sized objects, has been measured to be at least 10x that of water (Erdel 2015; Speil 2010), but for chromosome sized objects diffusing through a dense polymer melt of other chromatin, we would certainly expect the viscosity to be several orders of magnitude higher.

Our simulations employ a spring constant for the link between nodes of magnitude 0.25 force units per distance unit which, given that one distance unit is equivalent to 100 nm and 0.15 force units (σ) are equivalent to 0.4 pN, gives a spring constant of 0.007 pN/nm. To see if this is reasonable, we calculate the Young’s modulus assuming that each link is a cylinder of length 100 nm and diameter 100 nm. With this geometry, a spring constant of 0.007 pN/nm for extension of the cylinder by an external force acting on the ends corresponds to a Young’s modulus of 100 Pa. This agrees well with experimental measurements on interphase chromatin for which a Young modulus of approximately 200 Pa has been reported (de Vries et al., 2007).

Our simulations used a range of unpairing probabilities (P_unpair) of 0.001 – 0.1. The frequency of unpairing in units of s^−1^ can be obtained from the probability of unpairing by noting that P_unpair is the frequency of unpairing events per timestep and then multiplying by a conversion factor of 10 timesteps per second (based on the assumption of 0.1 s per timestep discussed above). This gives us a range of unpairing frequencies of 0.01 - 1 s^−1^. In vitro kinetic measurements have found that the dissociate rate for RecA mediated DNA pairing is on the order of 0.1 s^−1^ (Bazemore et al., 1997), consistent with the range of unpairing frequencies we have employed.

We summarize the following conversion factors that relate parameters of the model to real units. One time step in the simulation corresponds to 0.1 s in actual time. One distance unit in the simulation corresponds to 100 nm in the physical cell. One force unit in the simulation corresponds to 2.7 pN.

Although parameter values were chosen based on order of magnitude estimates from yeast, it should be noted that the simulations reported here are not intended to accurately represent molecular details of an actual biological system, but rather to test the concept of telomere force-driven discrimination of pairing using a highly simplified model.

### Simulating competition between homologous and homeologous chromosomes

To simulate competition between homologous and homeologous regions, we initialize the simulation with a total of four chromosomes, numbered 1-4. Chromosomes 1 and 2 are homologous to each other and homeologous to chromosomes 3 and 4. Chromosomes 3 and 4 are homologous to each other and homeologous to chromosomes 1 and 2. We will refer to pairing of a homologous locus with its corresponding locus on the homologous chromosome as correct pairing. If a locus pairs with the corresponding locus on either of the two homeologs, this is referred to as incorrect pairing. Within the simulation, unpaired nodes at corresponding positions on homologous or homeologous chromosomes are switched to the paired state whenever they approach each other within a pre-set capture radius. The probability of pairing is assumed to be the same regardless of whether the pairing is correct or incorrect. At each time point, paired loci have the possibility to switch to an unpaired state, with a probability that is different for correct versus incorrect pairings. The probability of switching from the paired to unpaired states in any given iteration are specified by the variables P_unpair_correct and P_unpair_incorrect. Unless specified otherwise, P_unpair_correct is always 0.001.

### Simulating Zippering in the presence of constant telomere force

Simulations for Figures 4-6 started from a half-paired (amphitene) state in order to avoid having to simulate the initial pairing process. A chain of length 50 was generated, after which a second copy of the same chain was added to the system in which all nodes are in the same position in space as the corresponding nodes in the original chain. The first 25 nodes were then initialized to the paired state, and the second 25 nodes initialized to the unpaired state. The Brownian dynamics simulation was run for 2000 iterations with the probability of pairing and unpairing both set to zero, allowing the unpaired segments to relax subject to nuclear volume constraint. At this point, the probability of pairing was set to 1 for loci within the capture distance, and the telomere force was activated. In contrast to the pairing simulations, for this analysis the telomere force was only applied to the last node in each chain, and the forces were directed in opposite directions that are constant over time. This non-fluctuating force was used in order to directly analyze the ability of zippering to do work against an applied force.

Unless otherwise stated, these tug-of-war simulations all employed a nuclear radius of 50, which is roughly five times higher than our estimated nuclear radius for yeast, but which we use to ensure that nuclear radius did not limit the extent to which the telomeres could be pulled apart. This point is discussed in the text and analyzed in Figure 6. Also, we note that for these simulations, the telomere force is always exerted constantly in one direction, and not with a randomly fluctuating orientation. The persistence of this applied force means that the magnitude of the force is not directly comparable with that used in previous simulations of pairing selectivity described above.

### Calculating stall force of chromosome zippering

For each value of P_unpair, an initial range of values for the telomere force magnitude ϕ were tested, with results assessed in terms of how many nodes were paired at the end of the simulation, until values were found that bracketed the stall force, as judged by the fact that one value allows zippering and one causes unzippering. Once these bracketing values were found, binary search was implemented by testing a value of ϕ halfway between the current bracketing values, and retaining whichever pair of values continued to bracket the stall force. The process was concluded when a value was found in which the average number of paired nodes was in the range 24-26. Because the simulations began with half of the 50 nodes paired, final outcomes with half the nodes paired reflect a situation in which the zippered region neither elongated nor shortened, in other words, a stall of the zippering process.

## Results

### Processive zippering decreases discrimination of pairing

In our previous simulation of meiotic homolog pairing, we found that even substantially high probability of unpairing still allowed processive zippering (Marshall and Fung, 2016). Avidity effects create a situation in which even weak affinity is enough to get strong pairing, since many sites all act together. This result raised the possibility that weak or inexact homologous regions might be able to effectively compete with regions of perfect homology. We therefore augmented our simulation framework to represent a situation in which the cell contains four chromosomes, which we will denote a, b, c, and d (**Figure 1A**). In the figure, chromosomes a and b are homologs of each other, whereas c and d are homologs of each other and homeologs of a and b. Each chromosome is modeled as a Rouse polymer represented by a bead-spring chain (**Figure 1B**) of 50 loci, such that corresponding loci on homologous chromosomes will pair if they come within a specified capture radius (**Figure 1C**), and will also unpair with a spontaneous unpairing probability P_unpair_correct, as in our previously published model. To simulate competition, as would occur with homeologous chromosomes, we also allow corresponding loci on a to pair with corresponding loci on c or d (the same is also true for chromosome b pairing with c or d) and vice versa. But in this case, if the pairing is between loci on a or b with loci on c or c, dissociation occurs with a different probability P_unpair_incorrect, which we assume is higher than P_unpair_correct. It is important to note from the outset that each locus has one correct locus that it can pair with, but two incorrect loci. With these assumptions in place, chromosome movement and pairing is simulated as described in materials and methods (**Figure 1D**).

**Figure 1.**
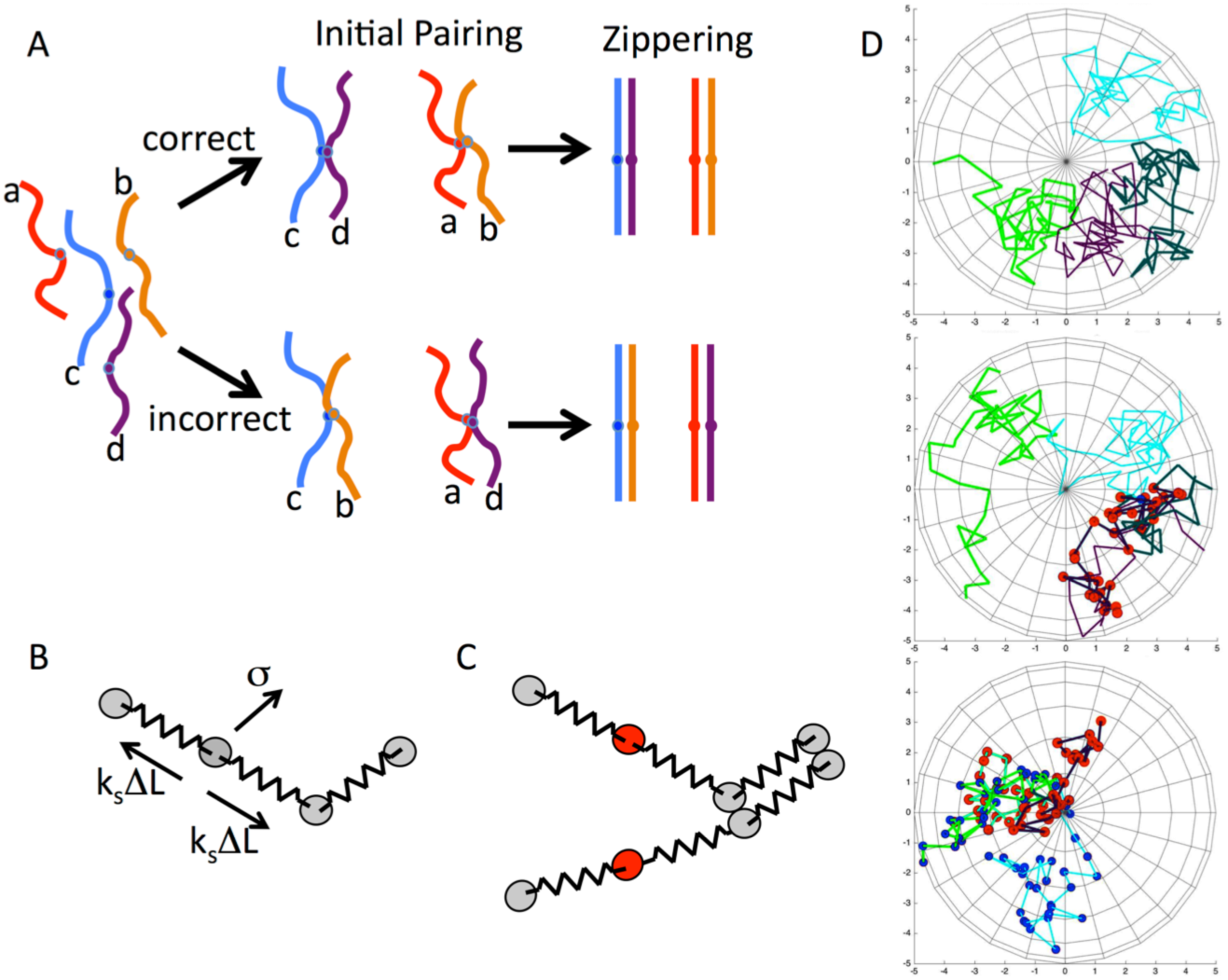
Simulating meiotic homolog pairing specificity. (**A**) Diagram depicts two pairs of chromosomes that share weak homology. Chromosomes labeled with lower case a and b are homologs of each other, and chromosomes c and d are homologs of each other. Chromosomes a and b are homeologs of c and d, so that they share weak homology that allows them to pair. Pairing is initiated at whatever locus happens to make first productive contact with either the homologous (top) or homeologous (bottom) chromosome. After initial pairing, zippering will cause the completion of pairing. Because of avidity effects, even weak pairing interactions can drive processive zippering, raising the question of how the scenario depicted on the bottom may be avoided. The color scheme depicted in this diagram will be used in subsequent figures to plot results of simulations. (**B**) Bead-spring model for chromosome dynamics as detailed in equation1. The chromosome is represented as a chain of beads connected by freely jointed springs. Each node experiences random thermal forces denoted by σ, as well as forces exerted by the two attached springs which are proportional to the stretch ΔL and the spring constant k_s_. The action of these forces is resisted by viscous drag. (**C**) Representation of homolog pairing within the bead-spring model framework. Nodes at corresponding locations along homologous chromosome chains are allowed to pair. Pairing of a node (shown in red) can only happen when it is closer to the corresponding homologous node by a fixed capture radius. (**D**) Images from a typical simulation run. Top panel depicts the chromosome arrangement at an early stage of the simulation (t=1000) before first pairing event. The four chromosomes are shown in different colors, with shades of green representing one set of homologs, and shades of blue the other. Middle panel (t=21000) showing initial incorrect pairing. Red spheres represent loci that have paired with the incorrect chromosome. Bottom panel (t=54000) showing a later stage of the simulation, during a time at which a combination of correct and incorrect pairing exists. Blue spheres indicate correctly paired loci, red spheres incorrectly paired loci.

The competition scenario outlined in **Figure 1A** was simulated under a range of unpairing probabilities, with a typical result depicted in **Figure 2A**. We depict the results by drawing each chromosome as a color bar, and then showing a second color bar alongside it that indicates which of the other three chromosomes it is paired with. Both simulations were run using a ten fold difference in unpairing probability between the correct and incorrect homologs. If all loci paired and unpaired independently, one would expect that roughly one sixth of the loci would be incorrectly paired, given that for each correct pairing partner there are two incorrect pairing partners. Clearly, however, large stretches of incorrect pairing can occur. To gain a more systematic view of the relation between selectivity at the level of unpairing rates versus at the level of overall pairing, simulations of pairing competition were run with a range of different unpairing rates for the incorrect associations. As plotted in **Figure 2B**, the result is that even with a more than ten-fold difference in affinity, there is predicted to be substantial degrees of mispairing on average.

**Figure 2.**
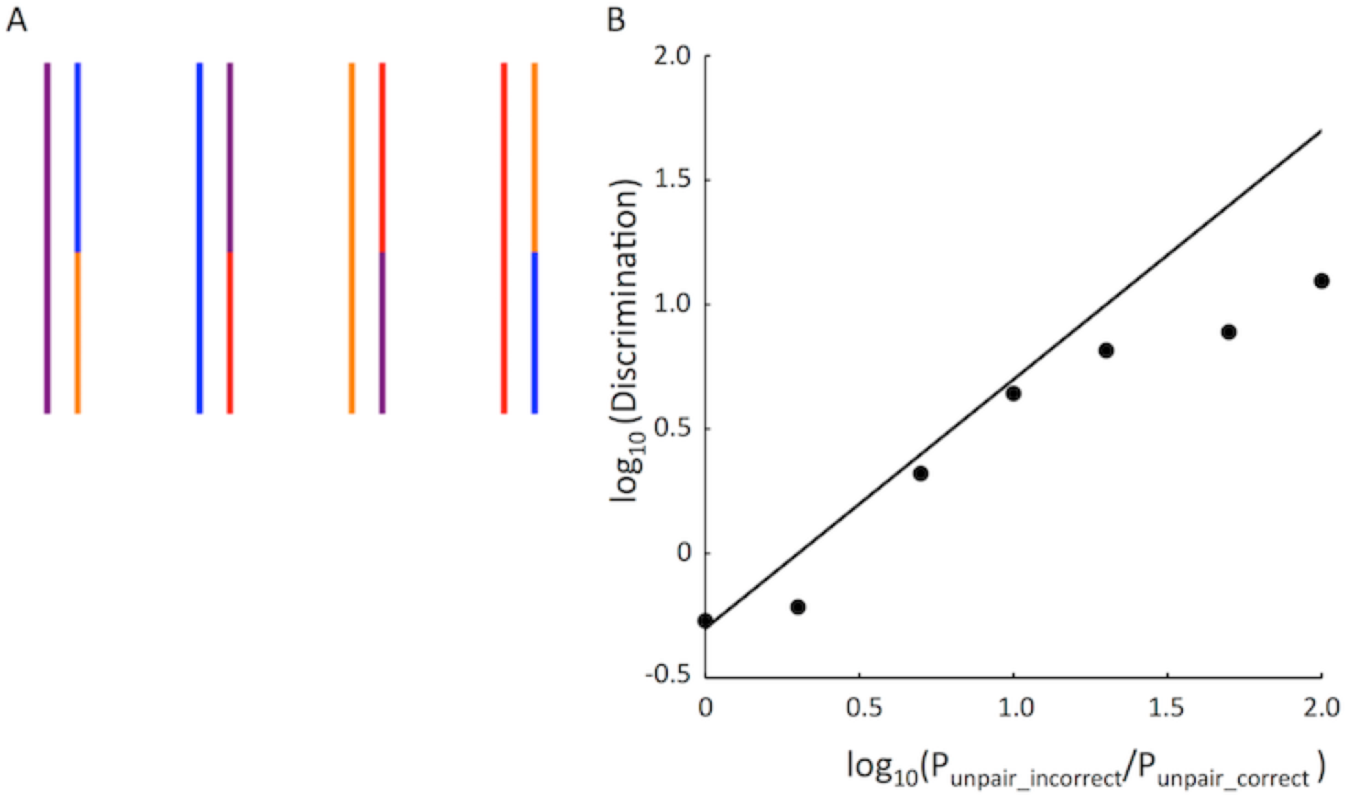
Zippering reduces pairing fidelity. (**A**) pairing outcome, plotted as in Figure 1A, for a typical simulation run with parameters P_unpair_correct 0.001 P_unpair_incorrect = 0.01. As illustrated by these two images, even with a ten-fold difference in affinity, incorrect pairing can be extensive. (**B**) Pairing discrimination, given as the ratio of correctly paired loci to incorrectly paired loci, plotted as a function of the unpairing probability (P_unpair) for incorrect versus correct loci. Simulations were run using P_unpair_correct of 0.001. (Circles) results of 100 simulations for each set of parameters. (Line) the expected discrimination if all loci paired and unpaired independently. Deviation of the simulation results below the line indicate that when loci are linked together into a polymer, discrimination is reduced, presumably due to avidity effects that prevent recently unpaired loci from diffusing apart.

### Active telomere forces can increase selectivity of pairing

The results of **Figure 2B** indicate that Brownian motion (in the form of thermally driven chromosome movement) is insufficient to overcome the avidity effect even for weak interactions. We can understand this effect by considering what happens when a pair of loci become unpaired. If they remain within the capture radius of each other during one time-step, they will immediately re-pair. Thus, unpairing can only happen if the just-unpaired loci diffuse apart from each other by a distance of more than the capture radius.

Perhaps active forces might be used to pull apart weakly bound regions to improve fidelity. In the framework just discussed, the extra force applied by the cytoskeleton would be able to drive recently unpaired loci far enough apart that they cannot immediately re-bind. On the other hand, it is not immediately obvious that application of such forces only to the telomeres would have an effect on the rest of the chromosome. To test whether active forces at the telomeres could enhance fidelity, we repeated the simulations of competition between chromosomes with high and low affinity in the presence of active randomly-directed forces applied to the telomeres. In these simulations, the magnitude of the active force ϕ was set to be 3.3 times the magnitude of the thermal random force, and switched direction with every timestep. The result of this simulation is shown in **Figure 3A**. For comparison, a line is also included which indicates the predicted discrimination if each locus paired and unpaired independently of the others in the absence of applied forces. When the ratio of unpairing probability is sufficiently close to 1, discrimination is similar in the presence of absence of the active random force. However, as the probability of unpairing for the incorrect loci increases, the discrimination becomes markedly better in the presence of active forces. Indeed for sufficiently large unpairing probability ratios, the active forces increase discrimination by two orders of magnitude. In vitro measurements of unpairing rate constants for RecA mediated DNA pairing between perfect homologs and homologs with mismatches showed that there was little discrimination for less than 10% mismatches but after that, increasing mismatches led to a difference in affinity of roughly 5-fold (Bazemore et al., 1997). Such a 5-fold difference in unpairing probability is where we begin to observe an effect caused by active forces in **Figure 3A**. We emphasize, as noted already, that the molecular mechanisms that mediate meiotic homology pairing remain controversial, hence RecA mediated DNA pairing may or may not be a relevant comparison, but at least it suggests that the range of differences in affinity that our model employs are at least biologically plausible.

**Figure 3.**
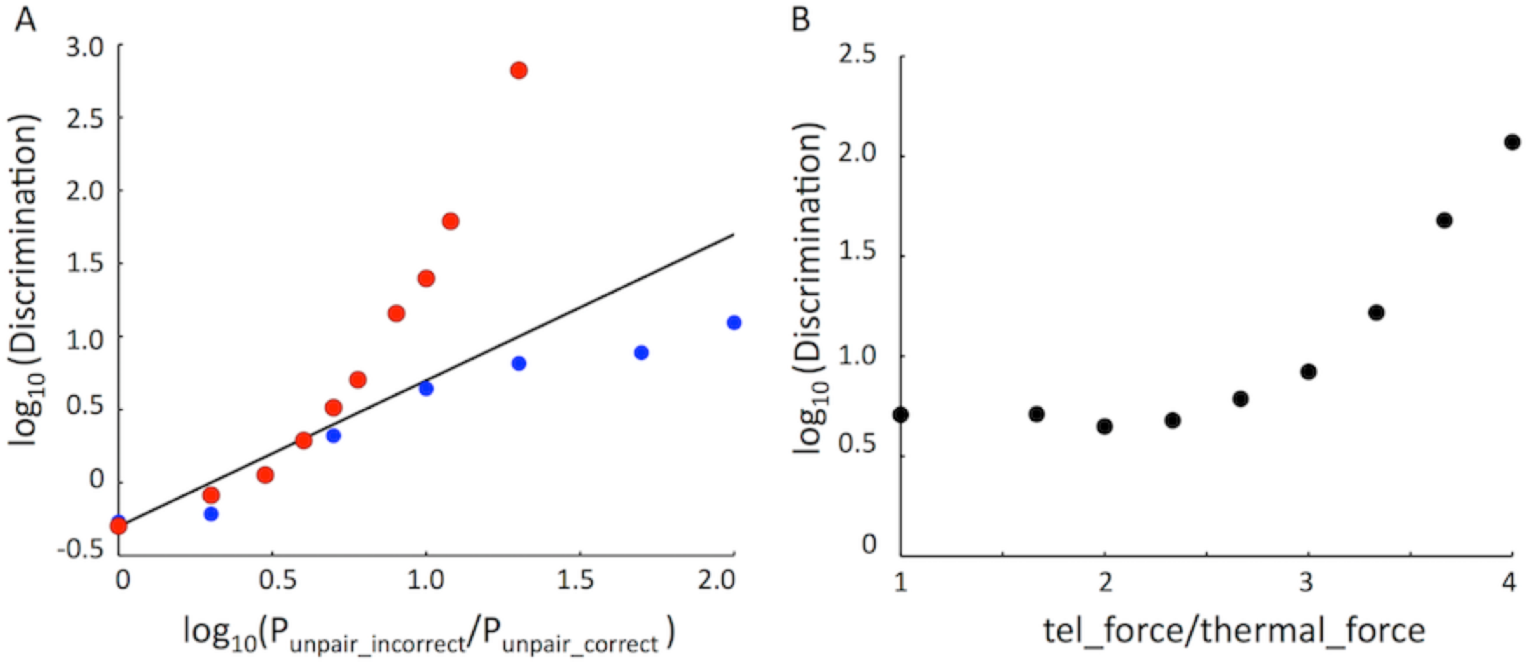
Active telomere forces can enhance selectivity of pairing. (**A**) Discrimination versus intrinsic selectivity of pairing for loci, in the presence versus absence of active telomere forces. Red markers indicate simulation results performed with telomere force ϕ = 0.5. Blue markers indicate simulation results in the absence of applied telomere force. Black line indicates expected discrimination if all loci pair and unpair independently. Deviation of red markers above the line indicates that active forces allow greater discrimination than possible by passive independent pairing and unpairing. Results are based on 50 independent simulations for each condition. (**B**) Discrimination versus magnitude of random telomere force. Force is expressed as the ratio of the applied telomere force ϕ to the simulated thermally-driven random forces acting at each node of the chain. Markers show results from 100 simulation runs for each condition. Simulations employed P_unpair of 0.001 for correct and 0.01 for incorrect loci.

We next asked how the increase in discrimination depends on the magnitude of the telomere random force (**Figure 3B**). Simulations were performed in which the unpairing probability was 10 fold higher for incorrect versus correct loci, and the magnitude of the telomere random force ϕ was swept through a range of values. When the magnitude of the telomere force was equal to the magnitude of the thermal random force, discrimination was the same as seen in the absence of any active force, as expected. Discrimination did not increase with small increases in the random force magnitude, but once the magnitude of the random telomere force exceeded the thermal force by a factor of two or more, discrimination improved as a monotonically increasing function of the magnitude of the force. Taken together, these analyses support the idea that random forces can, at least in principle, increase fidelity of meiotic pairing.

### A tug of war between telomere forces and a pairing-based Brownian ratchet

How does a mechanical property like the magnitude of a random force have an effect on a molecular property like discrimination of correct from incorrect loci? The processivity of zippering is suggestive of a mechanical system capable of force generation. Specifically, as the two homologous chromosomes zip together, the unpaired telomeres will be pulled closer. This large-scale mechanical behavior, which has the potential to perform work if the telomeres are pulled against an opposing force, is ultimately driven by the molecular-level interactions between homologous nodes in the chain. Zippering thus potentially creates a relationship between molecular recognition and chromosome mechanics.

We characterized the ability of zippering to generate force by simulating a condition in which the chromosomes start out fully paired along half their length, and then simulation pairing and unpairing in the presence of continuous force applied at the telomeres at one end of each chromosome (**Figure 4A**). In these simulations, forces are applied in a uniform direction at each telomere, so that the telomeres would tend to be pulled apart. There are two possible outcomes – either the force will drive unzippering of the region as the chromosome ends are pulled apart, or the zippering will pull the chromosome ends together despite the applied force.

**Figure 4.**
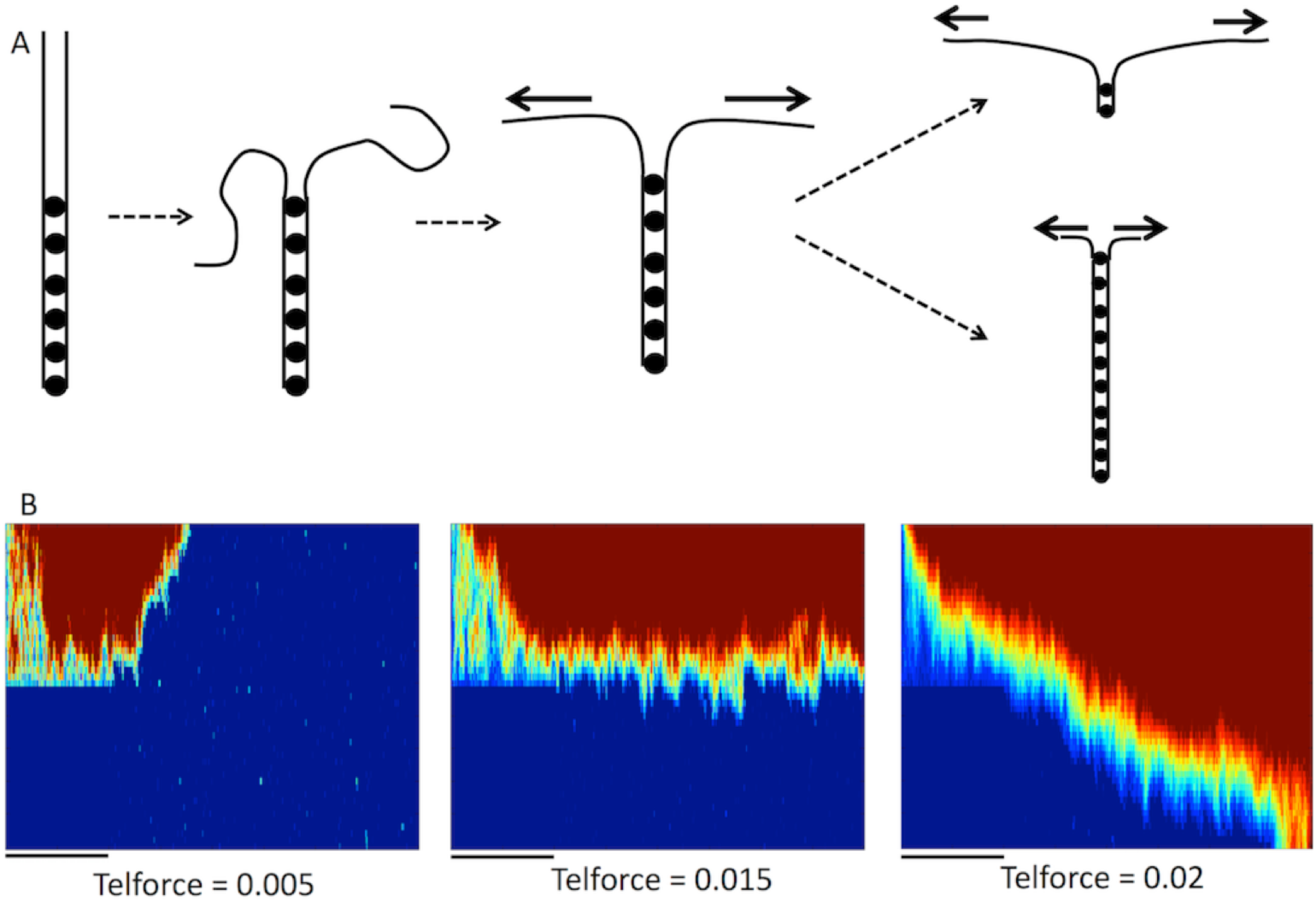
Mechanical competition (tug of war) between processive zippering and active telomere forces. (**A**). Simulation of tug of war. Two chromosomes, represented as beads connected by springs, are initialized into a state where half the nodes are paired (indicated by dots) and the other half is unpaired. The unpaired region is then allowed to relax under influence of Langevin random forces. Next, active forces are applied at the unpaired telomeres, and the simulation is run with this constant force applied in a uniform direction. As illustrated on the right, two possible outcomes are that either the telomere force is able to pull the chromosomes apart, leading to unzippering, or else the zippering can overcome the telomere pulling and drive further pairing beyond the initial extent of pairing. (**B**) Pairing kymographs showing results for three different magnitudes of telomere force. Black bars under each kymograph indicate the initial 2000 timepoints during which the unpaired region is allowed to stretch while the initial paired region is locked in the paired state. During these simulations, the telomere forces are applied in constant, opposite directions at the two telomeres. It is important to note that unlike previous simulations, the telomere force is applied in a constant direction, and does not fluctuate. All kymographs in this figure were generated using an unpairing probability of 0.01, with the different forces applied at the telomeres as indicated. All simulations for this figure were performed using a nuclear radius of 50.

Outcomes of these simulation results can be visualized in the form of pairing kymographs (Marshall and Fung 2016), in which different colors are used to represent the distance between homologous loci, with dark blue representing zero distance, and red representing the average distance between completely unpaired loci (**Figure 4B**). Shades of yellow indicate regions at intermediate distances. To generate a kymograph, the inter-homolog distance map at any point in time is plotted as a vertical color bar, with the top and bottom of the bar representing the two ends of the chromosomes. These color bars are then stacked left to right in temporal order. Pairing at any given locus is indicated by a switch from red to blue as one traverses the plot from left to right, while unpairing is represented as a switch from blue to red. Processive zippering is indicated by upwardly diagonal edges, while unzippering is indicated by downwardly directed diagonal edges. Each of these pairing kymographs illustrates a simulation of 8000 timepoints.

To perform the simulations, the two homologs were initially paired along their entire length, and then the first 25 nodes are set to the unpaired state while the second 25 nodes are locked in the paired state (P_unpair=0). The simulation was run for 2000 time points (indicated by black bars under each kymograph), giving the telomeres time to separate under the influence of the active forces. At t=2000, the unpairing probability for the second 25 nodes was set to a value of P_unpair = 0.01. The simulation was then run for an additional 6000 time points. Representative results are illustrated in **Figure 4B** for three different values of the telomere force magnitude ϕ. Because we did not allow the telomere forces to vary in direction, but rather applied a constant force at each telomere in opposing directions, the values of ϕ in these simulations should not be directly compared to those in the prior simulations, in which the direction of the forces fluctuated randomly between time points. The purpose of this simulation was not to simulate actual events during meiosis, but to test whether avidity-driven zippering could mechanically oppose a constant force.

As illustrated by the examples in **Figure 4B**, when the telomere force is weak, zippering still occurs. When the telomere force is sufficiently strong, the chromosomes become completely unzipped. At an intermediate force, a stall condition is reached in which neither zippering nor unzippering occur.

We can understand the results of **Figure 4** by imagining a tug of war between constant active force applied at the telomeres by the cytoskeleton, and an opposing force generated by processive zippering, which can be viewed as a Brownian ratchet (**Figure 5A**). To see how this ratchet might work, we note that each step of zippering requires the two unpaired nodes following the last paired nodes to pair. Once this happens, the paired region extends by one node, and the unpaired telomeres will be drawn slightly closer together. Pairing requires unpaired loci to move within a specified capture radius. In the Brownian ratchet model, once the unpaired segments are maximally stretched by the applied telomere forces, pairing of the next set of unpaired nodes happens when the end-to-end length of one or both of the unpaired segments undergoes a thermally-driven positive length fluctuation such that the distance Δx between the two unpaired nodes becomes less than the capture distance. The unpaired segments can be viewed as springs in such a picture. The spring constant and equilibrium end-to-end length for a freely jointed chain are both functions of the chain length, such that as the chain length increases, the spring constant decreases as the inverse of the chain length (Grosberg and Khokhlov, 1994). This spring-like behavior of the chains means that as the chain is stretched more and more by the telomere forces, the probability of thermal forces driving enough length fluctuation to pair the next set of nodes decreases, eventually making further pairing impossible.

**Figure 5.**
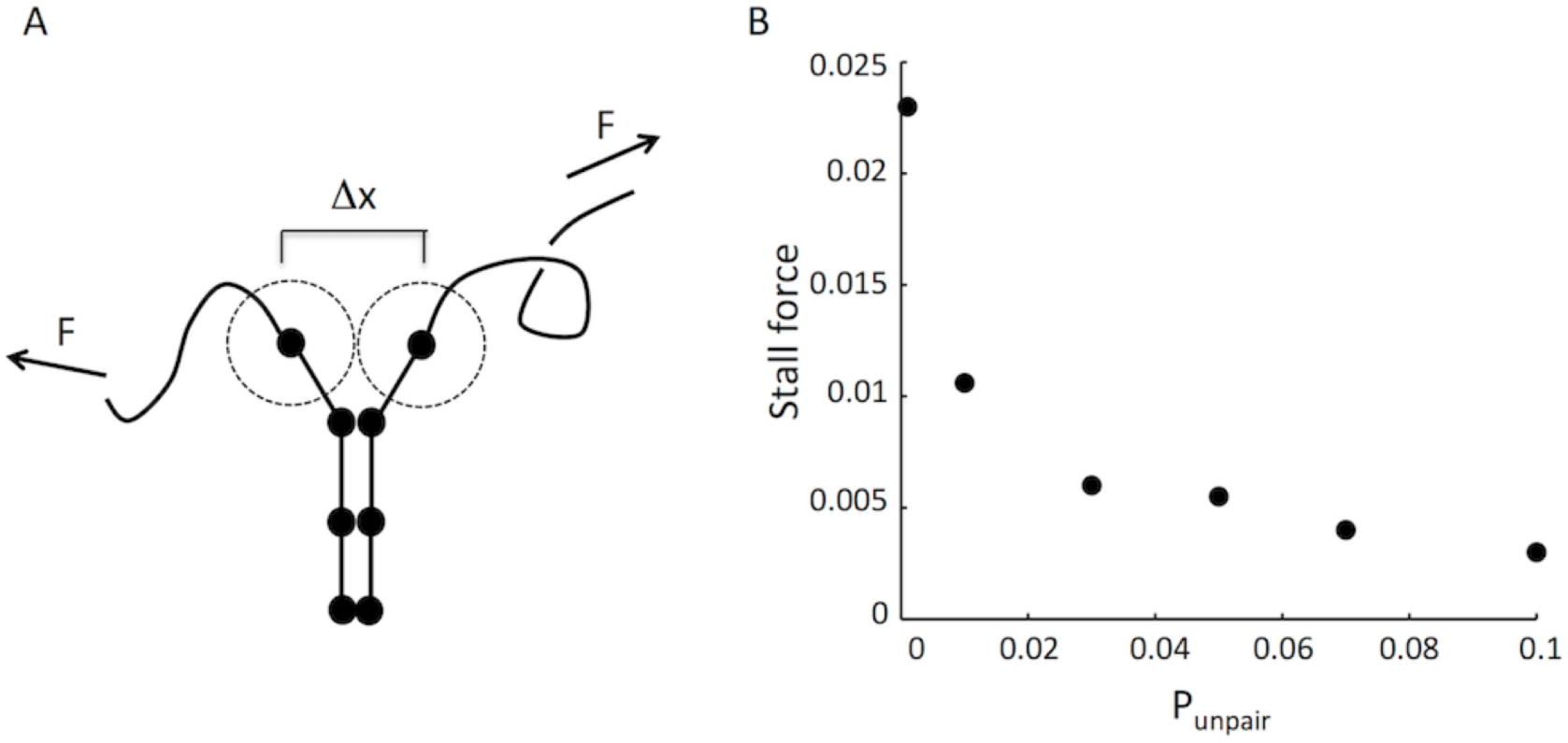
Brownian Ratchet model for force generation by zippering. (**A**) Telomere forces (arrows) tend to pull telomeres apart, which has the effect of exerting a force on the rest of the chromosome, leading to unzipping. However, thermal fluctuations in the length of the unzipped region will have a chance of allowing unpaired loci near the paired region to move within their respective capture radii (indicated by dotted circles). If this happens, pairing will occur and the zipped region will extend, despite the applied telomere force. The probability of loci moving the distance necessary to become paired, (Δx), will be a decreasing function of the magnitude of the telomere force. (**B**) Stall force versus unpairing probability. As discussed within materials and methods above, one force unit in the simulation corresponds to 2.7 pN, the range of forces given on the Y axis corresponds to a range of 0-0.07 pN.

In this tug of war scheme, the probability of zippering versus unzippering depends on the telomere force and the unpairing probability. A forward zippering step will take place if the first-passage time for the end-to-end lengths increasing is less than the dwell time between successive unpairing events. The smaller P_unpair is, the more chance there is that the unpaired segments will undergo a large enough end-to-end extension to allow capture of the next unpaired nodes. In contrast, if P_unpair is large, then the paired nodes at the end of the paired regions are expected to unpair before the next set of nodes are able to fluctuate within one capture radius of each other. Because the first passage time for the spring endpoints moving within a capture radius depends on the telomere force, we expect that for any given P_unpair, there should be a corresponding value of the telomere force ϕ at which the probability of a forward step equals the probability of a backward step. This represents a stall force for the zippering Brownian ratchet. We explored the stall force of this system by carrying out simulations at a range of P_unpair (**Figure 5B**). The stall force is seen to be a decreasing function of P_unpair. This result indicates that chromosomal regions that differ in their pairing affinity will have different stall forces. Given two different values of P_unpair for correct versus incorrect associations, it is thus possible to find a range of telomere forces which are able to stall or even reverse the zippering of weakly paired regions, while still permitting zippering of strongly pairing regions.

### Constraints due to nuclear diameter

The simulations of **Figure 4** and **5** were conducted using parameters in which the diameter of the nucleus is large relative to the relaxed length of the chromosomes, allowing the paired chromosomes to be fully pulled apart. The logic of choosing an initially large radius for the nucleus was that as the telomere force increases, the unpaired portion of the chromosome will stretch out more and more (**Figure 6A**), but cannot stretch to a length greater than the radius of the nucleus. If the nucleus is too small, then even large telomere forces would not be able to complete unzippering, simply because there would not be enough space for the unpaired regions of the chromosomes to stretch out (**Figure 6B**).

**Figure 6.**
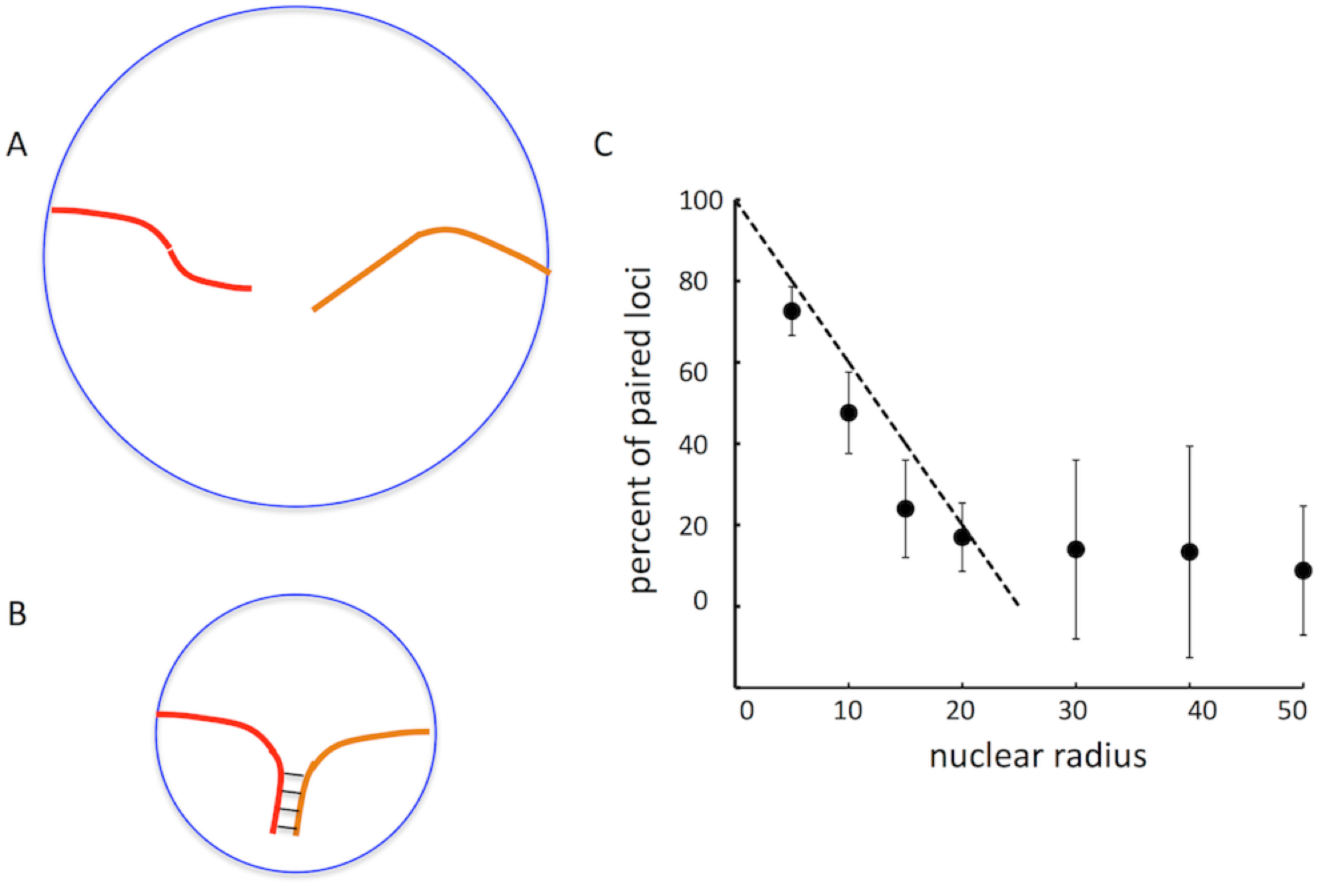
Extent of unzippering is constrained by nuclear envelope confinement. (**A**) For sufficiently large nuclei, strong telomere forces can in principle cause complete unpairing of chromosomes, potentially allowing homeologous or other incorrectly paired regions to be actively unpaired. (**B**) For small nuclei, whose radius is comparable to or less than the length of the chromosomes, even strong telomere forces cannot cause complete unpairing, because the maximum distance the telomeres can be pulled apart is the diameter of the cell. (**C**) Simulation results of tug of war implemented as a function of nuclear radius. Markers show percentage of paired loci (based on a maximum of 50 for complete pairing as in the tug-of-war simulations of Figure 4 and 5) after completion of simulation timecourse. Dashed line depicts the prediction for a simple model in which extent of unpairing is directly proportional to the nuclear radius.

We therefore explored the effect of nuclear radius on the tug of war simulations. We observed that the extent of unpairing achievable with strong telomere forces was not dependent on nuclear radius until the radius dropped below 25. Beyond this point, the number of unpaired nodes dropped proportionally with the radius (**Figure 6C**). This is the expected result given that the link length of the chromosome chain is 1 unit per link, such that the length of the initially unpaired segments would, if fully stretched, be 25 for each homolog. At maximal stretch, the two unpaired segments would thus fully span a nucleus of radius 25, and it would not be physically possible to pull them any further apart. For smaller radii, the unpaired segments could not fully stretch out unless zippering proceeded, resulting in a smaller number of unpaired nodes at the end of the simulation.

## Discussion

### Increased selectivity by randomly directed telomere forces

The processivity of homolog pairing speeds up pairing, but here we have shown that it also creates a problem for fidelity of the process, in that avidity effects allow even weakly paired loci to zip up, leading to potentially incorrect associations. **Figure 2** demonstrates that this effect causes the discrimination of correct from incorrect homologs to be less than expected if the individual loci were allowed to pair and unpair independently.

Our simulations (**Figure 3A**) indicate that random active forces, even when they are only applied at the telomeres, can enhance the discrimination in meiotic homology pairing. Interestingly, we found that active telomere forces were able to increase discrimination above the level expected by independent pairing/unpairing of loci. We propose that this increased discrimination can be explained by a tug of war model, in which zippering is mechanically opposed by the tendency for telomeres to be driven apart from each other. The stall force of zippering is expected to depend ion the off-rate of pairing, such that weakly paired regions would have a lower stall force. If the average force pulling the telomeres apart is greater than the stall force for incorrectly paired regions, but smaller than the stall force for correctly paired regions, the effect of the force would be to unzip any incorrect associations, while still allowing correct associations to proceed. In such a case, perfect discrimination should be possible. One point to note about this model is that our simulations of discrimination (**Figure 3**) employed randomly oriented, fluctuating telomere forces, while our simulations of the tug of war between zippering and telomere pulling (**Figure 4 and 5**) used constant telomere forces acting in a uniform direction. It is, however, the case that random forces acting on two objects (two telomeres, in this case) will tend to oppose the close apposition of those two objects, and will be able to drive them apart, at least over some range of distances.

The role of meiotic chromosome motion in unpairing was recently probed with chemical inhibitors to transiently arrest motion in fission yeast, followed by washout to restore motion, monitoring pairing during the process (Chacon et al., 2016). In that study, it was found that loci normally experienced fluctuating pairing states, such that a given locus would pair and unpair many times during the course of prophase. However, loci that happened to be paired when the motion was stopped, failed to unpair once the motion was resumed. It was inferred that the prolonged pairing created a pairing-locked state that could not be readily unpaired. Based on our simulation results, we suggest an alternative interpretation. Transient arrest of motion would allow paired loci to initiate zippering, which would, if allowed to progress long enough, lock the sites together such that even when motion resumed, they could not be unpaired. This would explain the lack of unpairing after transient motion arrest. Moreover, our results suggest that processive zippering in the absence of motion would lead to erroneous pairing with heterologous or homeologous chromosomes, which might then produce recombination intermediates that could not be properly resolved.

We note that active forces are likely to play multiple roles, with active unzippering being just one of them. In organisms like C. elegans, in which pairing is determined by highly specific pairing centers whose interactions are mediated by specific proteins, the initial interactions presumably have sufficient fidelity that extensive zippering of incorrect regions is unlikely to be an issue. For species with abundant repeat sequences widely distributed in the genome, incorrect zippering would be a serious problem. We therefore suspect that different species will vary substantially in the degree to which active motion contributes to pairing fidelity. Such considerations lead us to expect that experimental outcomes of abrogating telomere motion may vary widely between species.

### Comparing simulation results with persistent versus randomly varying forces

In the results above, we first showed that randomly fluctuating forces acting at the telomeres were able to enhance pairing selectivity (**Figure 3**). We then argued that telomeric forces are engaged in a tug of war with chromosome zippering such that forces exist that can unpair low affinity interactions but not high affinity interactions (**Figures 4 and 5**). However, the tug of war simulations employed forces acting in constant, opposite directions, so as to pull the chromosomes apart. This choice was made in order to directly probe the mechanical aspect of unpairing. But how then do the results of the tug of war analysis, in which directionally persistent forces are applied to push the telomeres apart, relate to the analysis of **Figure 3**, in which randomly fluctuating forces were used, whose direction was unbiased with respect to the relative position of the two telomeres? If we consider two objects, each undergoing independent random walks, the distance between them will tend to increase proportionally to the square root of elapsed time. This tendency to constantly increase the distance can be viewed as creating an effective force pulling the objects apart, albeit a force that varies as a function of the distanced between the objects. Given that the two telomeres are constrained to lie on the nuclear surface, there will be an average distance between them after which there will no longer be a consistent tendency for them to pull further apart. But if the telomeres were to be come very close, as in the final stages of zippering, then most random fluctuations would in fact tend to increase the distance between the telomeres, just as would be the case with a persistent directed force.

### Alternative mechanisms for force-based selectivity

The simplified model implemented here assumes that homologous loci dissociate with a constant probability that does not depend on the applied force. Because of this assumption, we explain the enhancement of discrimination in terms of telomeric force acting in opposition to zippering by slowing or stalling the continued pairing of unpaired sites, thus giving already-paired sites a chance to dissociate before they are locked in by zippering. However, an alternative hypothesis is that pulling forces exerted between chromosomes might increase the dissociation rate for weakly homologous loci, by providing enough energy to overcome the energetic barrier for dissociation. Such an alternative model would be equivalent to the idea that forces provide sufficient additional energy at the B cell receptor to overcome the energetic barrier for dissociation of lower affinity antigens (Natkanski et al., 2013). Single molecule experiments in which DNA strands are pulled apart by optical tweezers (Danilowicz 2003) have shown that the rate constant of dissociation depends not only on the base pair composition and the degree of nucleotide mismatches between the two strands, but also on the applied force (Hatch 2007). In meiosis, we predict that physical pulling applied to the chromosomes should increase the rate of unpairing, with the applied pulling force more likely to pull apart incorrect regions that lack extensive homology and thus cannot anneal as strongly. In this case, active forces would increase meiotic fidelity by catalyzing reversals of transient pairing. Our current simulation, in which the unpairing probability is assumed constant, argues that such selective unpairing is not required, but we cannot rule out that it might also make an additional contribution to fidelity in vivo. It is intuitively obvious that if a force-dependent dissociation is introduced, this will enhance the ability of forces to promote pairing fidelity, thus our central conclusions would still hold.

### Implications for nuclear size and elasticity

As shown in **Figure 6**, if the nucleus is not sufficiently large, active unzippering can be impeded simply by virtue of the fact that the telomeres cannot get farther apart than the diameter of the nucleus. In light of this observation, it is worth noting that nuclei tend to be significantly larger in meiotic cells than in mitotic cells of the same species. This enlargement is usually assumed to be required to support high levels of protein synthesis during egg formation. Based on the results of **Figure 6**, however, we hypothesize that the large size of meiotic nuclei may, at least in part, contribute to pairing fidelity by allowing more extensive unzipping. There is, however, another solution to the constraint imposed by nuclear size. If the nuclear envelope is sufficiently stretchable, then it is possible, at least in theory, for active forces to pull telomeres apart by a distance greater than the equilibrium nuclear diameter by deforming the nuclear envelope. This would be manifest as chromosomes that extend beyond the primary mass of chromosomes inside the nucleus. Such “maverick” chromosomes have been directly visualized in budding yeast meiosis (Scherthan et al, 2007). The fact that telomere forces appear to act over such long distances that chromosomes move apart from the rest of the nuclear mass is consistent with our hypothesis that these forces are serving to unzip incorrectly paired regions. In contrast, the existence of maverick chromosomes is less consistent with the idea that the primary role of these forces is to promote pairing by increasing the frequency of chromosome collisions, since the maverick chromosomes are actually pulled away from the other chromosomes. In any case, it is clearly important to develop experimental strategies to explore the interplay of nuclear radius and maverick chromosome behavior on different aspects of meiotic pairing selectivity and kinetics.

## Acknowledgments

This work was supported by NIH grant R01GM116895 (J.C.F.) and by NSF grant DBI-1548297 (W.F.M. and J.C.F.).

